## Supplementary material for "The secret life (cycle) of temperate bacteriophages"

**This PDF file includes:**

Figs. S1 to S8
Tables S1 to S4

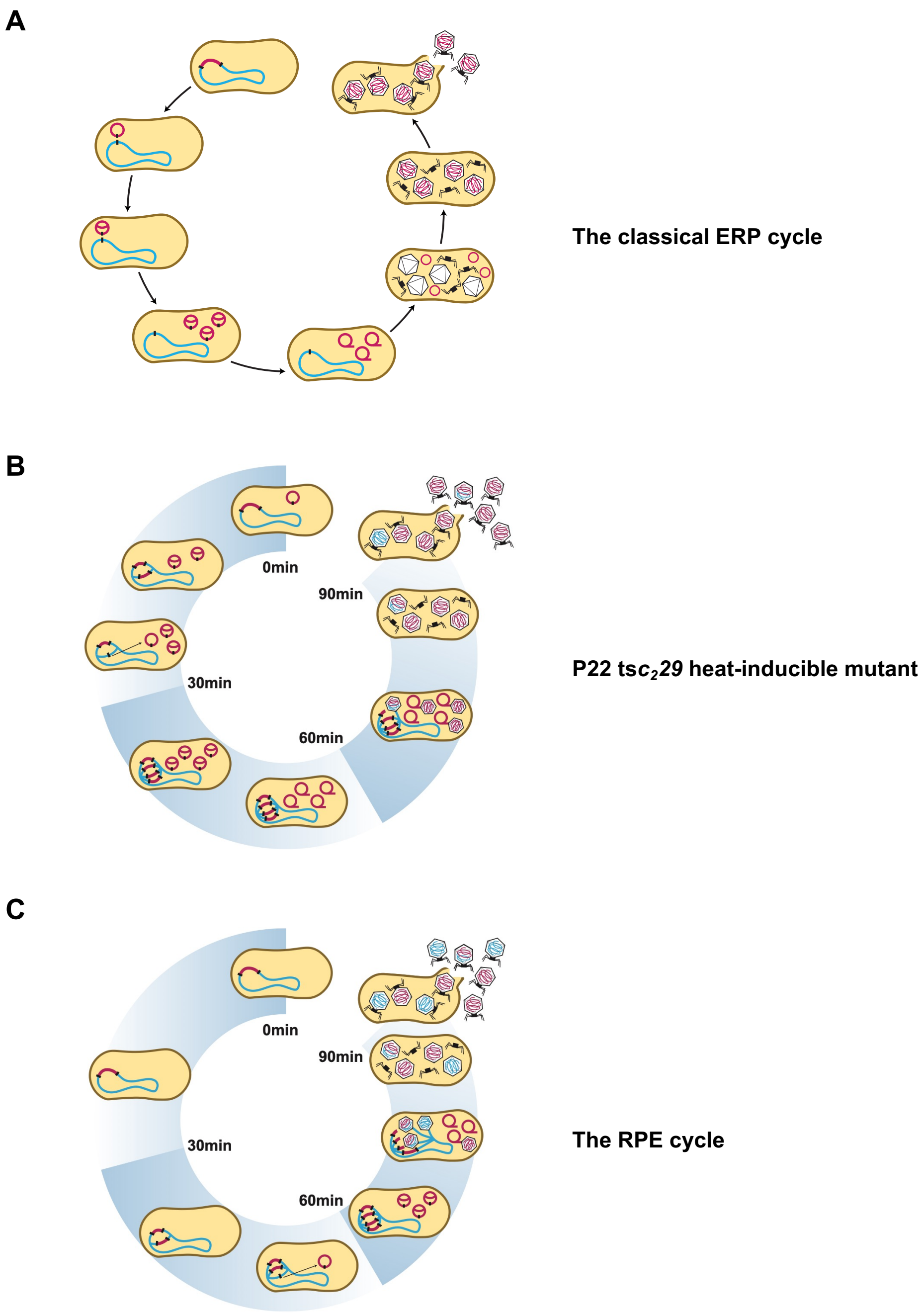


**Figure S1. The life cycle of temperate prophages after induction. (A)** Representation of the classical ERP model for prophage induction. **(B)** The P22 ts$c_2$*29* heat-inducible mutant follows the ERP cycle. **(C)** Representation of the RPE model. In red, phage genome; in blue, chromosomal DNA.

**A**

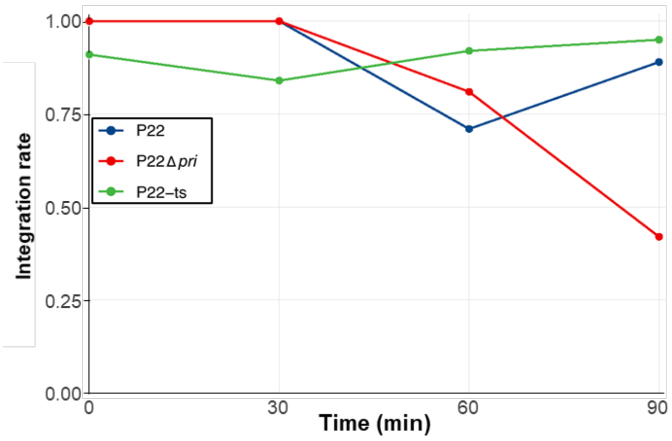


**B**

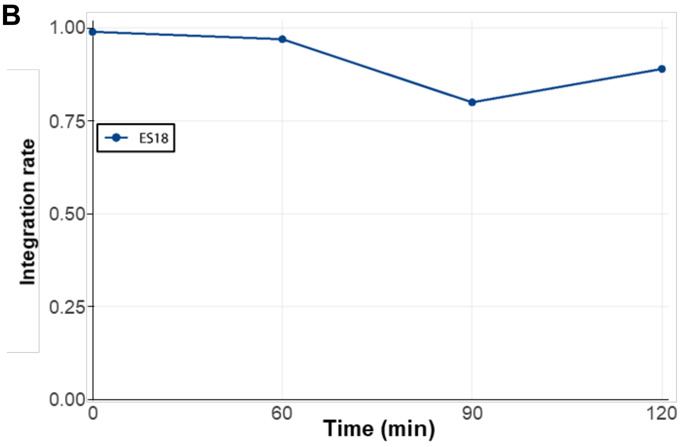


**Figure S2. Prophages excise late from the bacterial chromosome.** To gain insight into the timing of phage excision, the proportion of prophages integrated within the bacterial chromosome was measured over the course of 120 mins following induction (with MC or temperature). Integrated phages were measured as reads spanning the chromosome-*att*L phage region. The proportion of integrated phage at each timepoint was calculated by dividing the number of chromosome-*att*L reads by the total number of reads obtained from the sample (sum of chromosome-attL phage region reads plus reads spanning an unlinked chromosome-chromosome region), for phages P22, P22 Δ*pri* and P22 ts$c_2$29 **(A),** and for ES18 **(B)**.

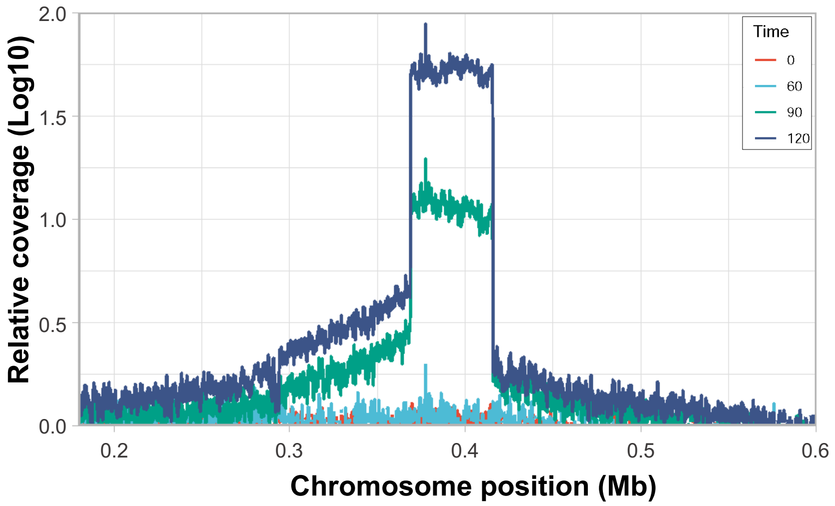


**Figure S3. Phage ES18 replicates *in situ* before excision.**
Relative abundance of phage genomic DNA and the chromosomal regions proximal to the integration for ES18 is represented. Samples were analyzed at 0 (red), 60 (cyan), 90 (green), and 120 min (blue) after induction (with mitomycin). Relative coverage is the DNA relative to the average bacterial genomic coverage (excluding phage).

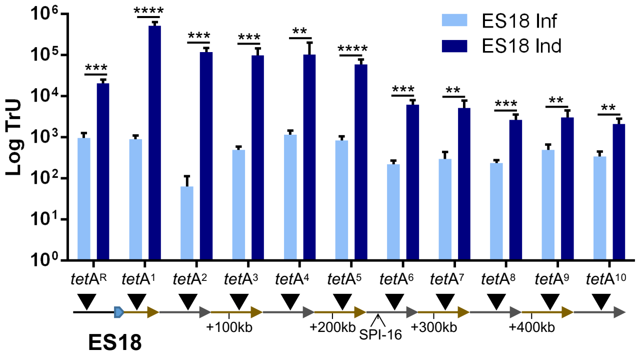


**Figure S4. ES18 engages in lateral transduction.** The transfer of *tet*A markers located upstream (*tet*AR) or downstream of the P22 *att*B site, in ten successive capsid headfuls (*tet*A"n"), was tested after ES18 induction or ES18 infection. Transduction units (TrU) per milliliter were normalized by PFU per milliliter and represented as the log TrU of an average phage titre (1 x $10^9$ PFU). Error bars indicate standard deviation. For all panels, values are means (n = 3 independent samples). An unpaired t test was performed to compare mean differences of infection and induction in each marker. Adjusted p values were as follows: *ns*>0.05; *p≤0.05; **p≤0.01; ***p≤0.001; ****p≤0.0001. The exact statistical values for each of the conditions tested are listed in Table S1.

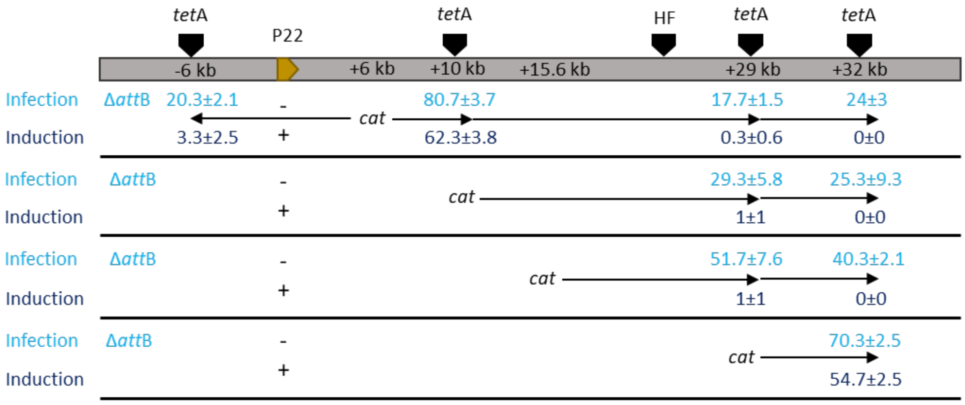


**Figure S5. Lateral transduction occurs by the headful mechanism.** Co-transduction frequencies for strains containing both *cat* and *tet*A markers at varying distances apart. Lysates were tested for co-transduction of both markers by initially selecting for the TetA marker, with transductants subsequently scored for the Cat marker. At least one hundred TetA transductants were tested from infection (cyan) or induction (blue) and the results are represented as a percentage (Cat/TetA) x 100%. The *att*B for P22 is highlighted in orange and the headful limit (HF) is indicated. The + indicates a P22 lysogen (induction, LT), while the - indicates a non-lysogenic strain (infection, GT). The means and standard deviations from three independent experiments are presented (*n*=3). Our results indicate that for lysates generated by P22 infection, all co-transduction frequencies were inversely proportional to their distance apart, even if they were located in two different headfuls, indicating that DNA packaging initiated at random sites. However, when we tested lysates generated by SOS induction, co-transduction was only observed for markers within a headful, indicating that packaging had primarily initiated from the bona fide *pac* site, and confirming that P22-mediated LT uses the headful mechanism for packaging.

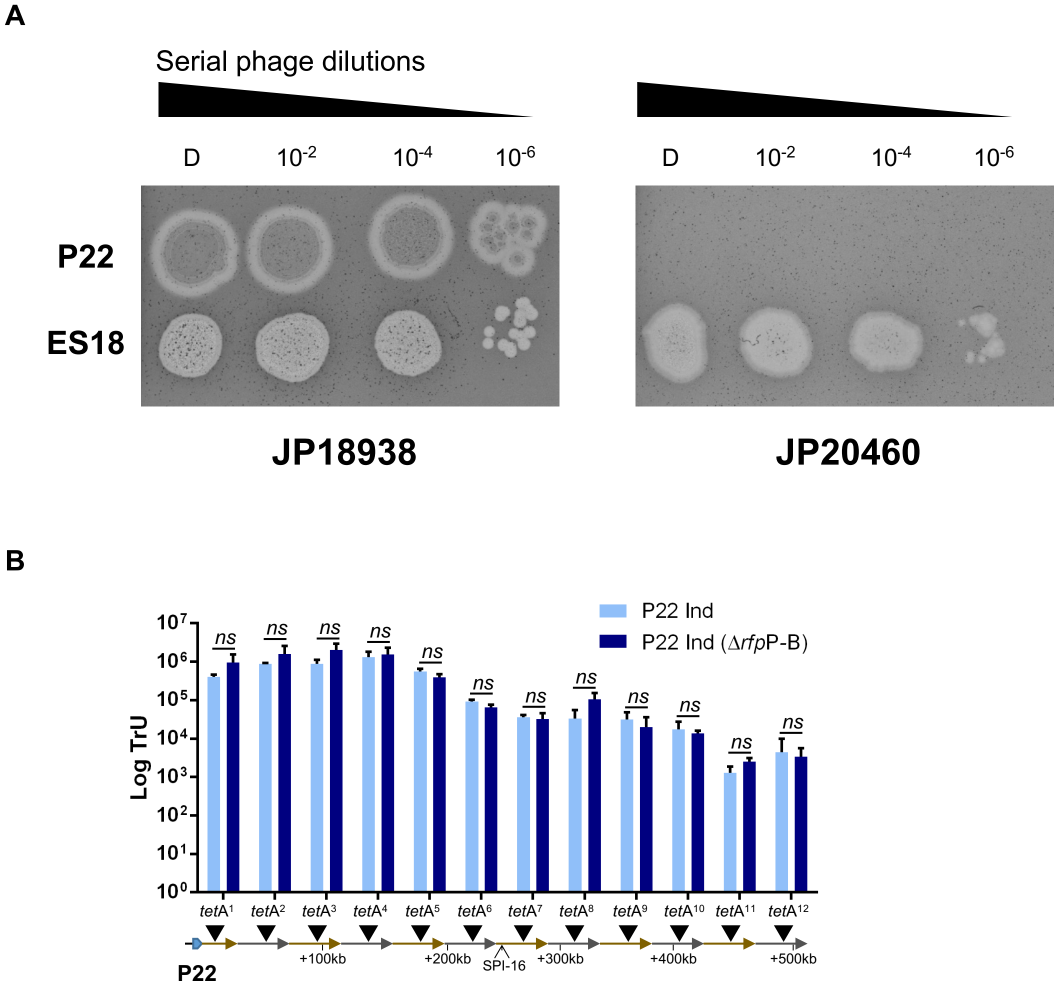


**Figure S6. Phage P22 engages in lateral transduction in an infection-resistant host.** To rule out the possibility that P22 phages released early after SOS induction could superinfect the remaining cells to initiate packaging from the resident prophage genome, we deleted the *rfp*P-B genes that encode the P22 receptor and tested this strain for LT. **(A)** Deletion of *rfp*P-B renders LT2 insensitive to P22 infection. Serial dilutions of phages P22 and ES18 lysates were spotted onto lawns of non-lysogenic LT2 and non-lysogenic LT2 *ΔrfpP-B* strains. The presence of plaques indicates successful phage infection and replication via the lytic cycle. **(B)** The transfer of *tet*A markers located downstream of the P22 *att*B site, in twelve successive capsid headfuls (*tet*A"n"), was tested after P22 induction in either wild type LT or LT lacking the phage P22 receptor (Δ*rfp*P-B). Transduction units (TrU) per milliliter were normalized by PFU per milliliter and represented as the log TrU of an average phage titre (1 x $10^9$ PFU). Error bars indicate standard deviation. For all panels, values are means (n = 3 independent samples). An unpaired t test was performed to compare mean differences of infection and induction in each marker. Adjusted p values were as follows: *ns*>0.05; *p≤0.05; **p≤0.01; ***p≤0.001; ****p≤0.0001. The exact statistical values for each of the conditions tested are listed in Table S1.

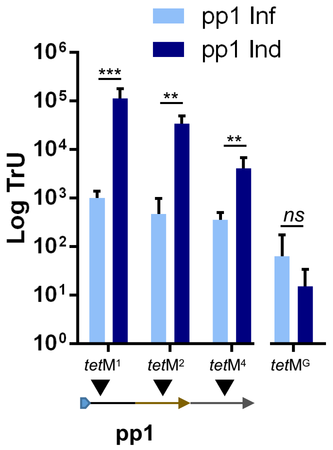


**Figure S7. Enterococcal phage pp1 engages in lateral transduction.** The ability of phage pp1 to mobilise *tet*M markers located downstream of the pp1 *att*B site in successive capsid headfuls (*tet*M"n"), or from non-phage associated regions of the chromosome (*tet*MG), was tested following induction and infection. Transduction units (TrU) per milliliter were normalized by PFU per milliliter and represented as the log TrU of an average phage titre (1 x $10^9$ PFU). Error bars indicate standard deviation. For all panels, values are means (n = 3 independent samples). An unpaired t test was performed to compare mean differences of infection and induction in each marker. Adjusted p values were as follows: *ns*>0.05; *p≤0.05; **p≤0.01; ***p≤0.001; ****p≤0.0001. The exact statistical values for each of the conditions tested are listed in Table S1.

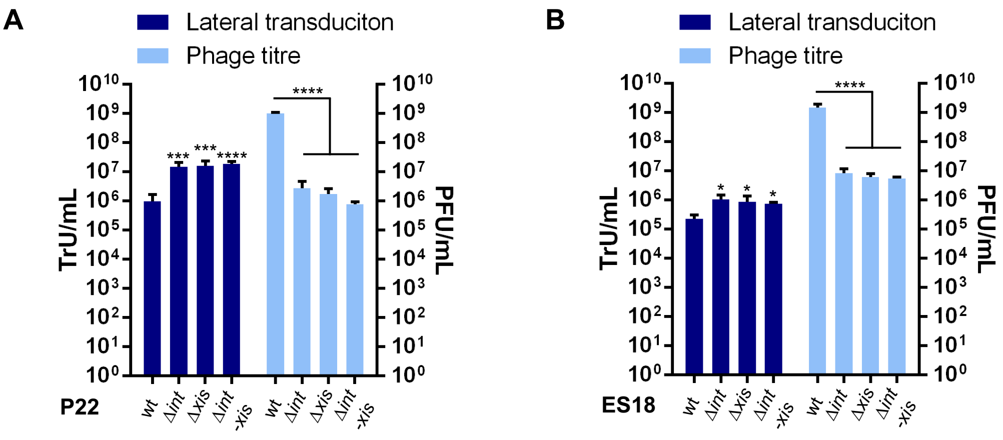


**Figure S8. Effect of phage mutations on lateral transduction and phage formation. (A)** P22 or P22-derivative (Δ*int*, Δ*xis* or Δ*int-xis*) prophages were mitomycin C induced and tested for their ability to transduce a *tet*A marker (TrU/ml) and produce viable phage particles (PFU/ml). **(B)** The same experiment was performed using phage ES18 or ES18-derivates (Δ*int*, Δ*xis* or Δ*int-xis*). The means and standard deviations from three independent experiments are presented ($n$=3). A one-way ANOVA with Dunnett's multiple comparisons test was performed to compare mean differences between time wt and derivatives phages. Adjusted p values were as follows: *ns*>0.05; *p≤0.05; **p≤0.01; ***p≤0.001; ****p≤0.0001. The exact statistical values for each of the conditions tested are listed in Table S1.

**Table S1. Statistical values.**

| **Figure 3A** | ***p* value** |
|---|---|
| HDR | 0.0009 |
| HD+1 | < 0.0001 |
| HD+2 | < 0.0001 |
| HD+3 | 0.0003 |
| HD+4 | < 0.0001 |
| HD+5 | < 0.0001 |
| HD+6 | 0.0002 |
| HD+7 | 0.0006 |
| HD+8 | 0.0006 |
| HD+9 | 0.0024 |
| HD+10 | 0.0009 |
| HD+11 | 0.0029 |
| HD+12 | 0.0266 |
| **Figure 3B** | ***p* value** |
| HD+1 | 0.0013 |
| HD+2 | 0.0009 |

| **Figure 5. Lateral transfer** | ***p* value** |
|---|---|
| 0-30 | 0.9942 |
| 0-60 | 0.0442 |
| 0-90 | < 0.0001 |
| 0-120 | < 0.0001 |
| 0-150 | < 0.0001 |
| 0-180 | < 0.0001 |
| 0-210 | < 0.0001 |
| 0-240 | < 0.0001 |
| **Figure 5. Phage titre** | ***p* value** |
| 0-30 | 0.9974 |
| 0-60 | 0.9892 |
| 0-90 | 0.8971 |
| 0-120 | 0.1527 |
| 0-150 | 0.0022 |
| 0-180 | < 0.0001 |
| 0-210 | < 0.0001 |
| 0-240 | < 0.0001 |

| Figure S4 | *p* value |
|---|---|
| HDR | 0.0002 |
| HD+1 | < 0.0001 |
| HD+2 | 0.0002 |
| HD+3 | 0,0001 |
| HD+4 | 0.0014 |
| HD+5 | < 0.0001 |
| HD+6 | 0.0001 |
| HD+7 | 0.0023 |
| HD+8 | 0.0005 |
| HD+9 | 0.0081 |
| HD+10 | 0.0041 |

| Figure S6 | *p* value |
|---|---|
| HD-1 | 0.0544 |
| HD+1 | 0.1756 |
| HD+2 | 0.2912 |
| HD+3 | 0.0739 |
| HD+4 | 0.843 |
| HD+5 | 0.1271 |
| HD+6 | 0.4411 |
| HD+7 | 0.5811 |
| HD+8 | 0.0631 |
| HD+9 | 0.6071 |
| HD+10 | 0.9967 |
| HD+11 | 0.1808 |
| HD+12 | 0.8182 |

| Figure S7 | *p* value |
|---|---|
| HD+1 | 0.0008 |
| HD+2 | 0.0023 |
| HD+4 | 0.0082 |
| *tet*$M^G$ | 0.9364 |

| Figure S8. P22 LT | *p* value |
|---|---|
| wt-Δ*int* | 0.0003 |
| wt-Δ*xis* | 0.0002 |
| wt-Δ*int-xis* | 0,0002 |
| **Figure S8. P22 titre** | ***p* value** |
| wt-Δ*int* | < 0.0001 |
| wt-Δ*xis* | < 0.0001 |
| wt-Δ*int-xis* | < 0.0001 |

| Figure S8. ES18 LT | *p* value |
|---|---|
| wt-Δ*int* | 0.0136 |
| wt-Δ*xis* | 0.0335 |
| wt-Δ*int-xis* | 0.0434 |
| **Figure S8. ES18 titre** | ***p* value** |
| wt-Δ*int* | < 0.0001 |
| wt-Δ*xis* | < 0.0001 |
| wt-Δ*int-xis* | < 0.0001 |

**Table S2. Strains used in this study.**

| Strain | Description | Reference |
|---|---|---|
| *E. coli* DC10B | K-12-derivative cloning strain; *dam*$^+$ *Ddcm DhsdRMS endA1 recA1* | {Monk:2012hc} |
| VE18590 | *E. faecalis* V583 derivative. Non-lysogenic, EfCIV583-negative. | (*39*) |
| VE18562 | *E. faecalis* V583 derivative. fp1 lysogen, EfCIV583-negative. | Matos:2013ca} |
| JP18938 | LT2 ΔFels-1 ΔGifsy-2 ΔGifsy-1 ΔFels-2 | This work |
| JP22117 | SV1208 ΔFels-1 | This work |
| JP18983 | JP18938 P22 lysogen | This work |
| JP18985 | JP18938 ES18 lysogen | This work |
| JP22119 | JP22117 P22 ts$c_2$*29* lysogen | This work |
| JP20460 | JP18938 Δ*rfb*P-*rfb*B::*Km*R | This work |
| JP20590 | JP18938 P22 lysogen Δ*int* | This work |
| JP20591 | JP18938 P22 lysogen Δ*xis* | This work |
| JP20592 | JP18938 P22 lysogen Δ*int-xis* | This work |
| JP20593 | JP18938 P22 lysogen Δ*orf12* | This work |
| JP20595 | JP18938 ES18 lysogen Δ*int* | This work |
| JP20596 | JP18938 ES18 lysogen Δ*xis* | This work |
| JP20597 | JP18938 ES18 lysogen Δ*int-xis* | This work |
| JP19020 | JP18938 Δ*att*B P22 | This work |
| JP22118 | JP22117 Δ*att*B P22 | This work |
| JP19086 | JP19020 *gpt*ΩTetA (6 kb upstream P22 *att*B) | This work |
| JP19087 | JP19020 STM0331ΩTetA (6 kb downstream P22 *att*B, HF1) | This work |
| JP19088 | JP19020 *prp*RΩTetA (47 kb downstream P22 *att*B, HF2) | This work |
| JP19089 | JP19020 *mal*ZΩTetA (88 kb downstream P22 *att*B, HF3) | This work |
| JP19090 | JP19020 *yaj*GΩTetA (130 kb downstream P22 *att*B, HF4) | This work |
| JP19091 | JP19020 *yba*NΩTetA (170 kb downstream P22 *att*B, HF5) | This work |
| JP19092 | JP19020 *ybb*VΩTetA (213 kb downstream P22 *att*B, HF6) | This work |
| JP19093 | JP19020 STM0557ΩTetA (246 kb downstream P22 *att*B, HF7) | This work |
| JP19094 | JP19020 *ahp*FΩTetA (304 kb downstream P22 *att*B, HF8) | This work |
| JP19095 | JP19020 *ybe*LΩTetA (348 kb downstream P22 *att*B, HF9) | This work |
| JP19096 | JP19020 STM0699ΩTetA (392 kb downstream P22 *att*B, HF10) | This work |
| JP19097 | JP19020 *nei*ΩTetA (425 kb downstream P22 *att*B, HF11) | This work |
| JP19098 | JP19020 *mod*FΩTetA (474 kb downstream P22 *att*B, HF12) | This work |
| JP20469 | JP18983 *gpt*ΩTetA (6 kb upstream P22 *att*B) | This work |
| JP20470 | JP18983 STM0331ΩTetA (6 kb downstream P22 *att*B, HF1) | This work |
| JP20471 | JP18983 *prp*RΩTetA (47 kb downstream P22 *att*B, HF2) | This work |
| JP20472 | JP18983 *mal*ZΩTetA (88 kb downstream P22 *att*B, HF3) | This work |
| JP20473 | JP18983 *yaj*GΩTetA (130 kb downstream P22 *att*B, HF4) | This work |
| JP20474 | JP18983 *yba*NΩTetA (170 kb downstream P22 *att*B, HF5) | This work |
| JP20475 | JP18983 *ybb*VΩTetA (213 kb downstream P22 *att*B, HF6) | This work |
| JP20476 | JP18983 STM0557ΩTetA (246 kb downstream P22 *att*B, HF7) | This work |
| JP20477 | JP18983 *ahp*FΩTetA (304 kb downstream P22 *att*B, HF8) | This work |
| JP20478 | JP18983 *ybe*LΩTetA (348 kb downstream P22 *att*B, HF9) | This work |
| JP20479 | JP18983 STM0699ΩTetA (392 kb downstream P22 *att*B, HF10) | This work |
| JP20480 | JP18983 *nei*ΩTetA (425 kb downstream P22 *att*B, HF11) | This work |
| JP20481 | JP18983 *mod*FΩTetA (474 kb downstream P22 *att*B, HF12) | This work |
| JP21928 | JP22118 *gpt*ΩTetA (6 kb upstream P22 *att*B) | This work |
| JP21929 | JP22118 *prp*RΩTetA (47 kb downstream P22 *att*B) | This work |
| JP21922 | JP22119 *gpt*ΩTetA (6 kb upstream P22 *att*B) | This work |
| JP21923 | JP22119 *prp*RΩTetA (47 kb downstream P22 *att*B) | This work |
| JP20482 | JP18985 *gpt*ΩTetA (6 kb upstream P22 *att*B) | This work |
| JP20483 | JP18985 STM0331ΩTetA (6 kb downstream P22 *att*B, HF1) | This work |
| JP20484 | JP18985 *prp*RΩTetA (47 kb downstream P22 *att*B, HF2) | This work |
| JP20485 | JP18985 *mal*ZΩTetA (88 kb downstream P22 *att*B, HF3) | This work |
| JP20487 | JP18985 *yba*NΩTetA (170 kb downstream P22 *att*B, HF4) | This work |
| JP20488 | JP18985 *ybb*VΩTetA (213 kb downstream P22 *att*B, HF5) | This work |
| JP20489 | JP18985 STM0557ΩTetA (246 kb downstream P22 *att*B, HF6) | This work |
| JP20490 | JP18985 *ahp*FΩTetA (304 kb downstream P22 *att*B, HF7) | This work |

| | | |
|---|---|---|
| JP20491 | JP18985 *ybe*LΩTetA (348 kb downstream P22 *att*B, HF8) | This work |
| JP20493 | JP18985 *nei*ΩTetA (425 kb downstream P22 *att*B, HF9) | This work |
| JP20494 | JP18985 *mod*FΩTetA (474 kb downstream P22 *att*B, HF10) | This work |
| JP19438 | JP20470 Δ*rfb*P-*rfb*B::*Km*R | This work |
| JP19439 | JP20471 Δ*rfb*P-*rfb*B::*Km*R | This work |
| JP19440 | JP20472 Δ*rfb*P-*rfb*B::*Km*R | This work |
| JP19441 | JP20473 Δ*rfb*P-*rfb*B::*Km*R | This work |
| JP19442 | JP20474 Δ*rfb*P-*rfb*B::*Km*R | This work |
| JP19443 | JP20475 Δ*rfb*P-*rfb*B::*Km*R | This work |
| JP19444 | JP20476 Δ*rfb*P-*rfb*B::*Km*R | This work |
| JP19445 | JP20477 Δ*rfb*P-*rfb*B::*Km*R | This work |
| JP19446 | JP20478 Δ*rfb*P-*rfb*B::*Km*R | This work |
| JP19447 | JP20479 Δ*rfb*P-*rfb*B::*Km*R | This work |
| JP19448 | JP20480 Δ*rfb*P-*rfb*B::*Km*R | This work |
| JP19449 | JP20481 Δ*rfb*P-*rfb*B::*Km*R | This work |
| JP21422 | JP19020 Cat marker 6 kb downstream P22 *att*B + TetA marker 6 kb upstream P22 *att*B | This work |
| JP21423 | JP19020 Cat marker 6 kb downstream P22 *att*B + TetA marker 10 kb up downstream P22 *att*B | This work |
| JP21424 | JP19020 Cat marker 6 kb downstream P22 *att*B + TetA marker 29 kb up downstream P22 *att*B | This work |
| JP21425 | JP19020 Cat marker 6 kb downstream P22 *att*B + TetA marker 32 kb up downstream P22 *att*B | This work |
| JP21426 | JP19020 Cat marker 10 kb downstream P22 *att*B + TetA marker 29 kb upstream P22 *att*B | This work |
| JP21427 | JP19020 Cat marker 10 kb downstream P22 *att*B + TetA marker 32 kb upstream P22 *att*B | This work |
| JP21428 | JP19020 Cat marker 15.6 kb downstream P22 *att*B + TetA marker 29 kb upstream P22 *att*B | This work |
| JP21429 | JP19020 Cat marker 15.6 kb downstream P22 *att*B + TetA marker 32 kb upstream P22 *att*B | This work |
| JP21430 | JP19020 Cat marker 29 kb downstream P22 *att*B + TetA marker 32 kb upstream P22 *att*B | This work |
| JP21431 | JP18983 Cat marker 6 kb downstream P22 *att*B + TetA marker 6 kb upstream P22 *att*B | This work |
| JP21432 | JP18983 Cat marker 6 kb downstream P22 *att*B + TetA marker 10 kb up downstream P22 *att*B | This work |
| JP21433 | JP18983 Cat marker 6 kb downstream P22 *att*B + TetA marker 29 kb up downstream P22 *att*B | This work |
| JP21434 | JP18983 Cat marker 6 kb downstream P22 *att*B + TetA marker 32 kb up downstream P22 *att*B | This work |
| JP21435 | JP18983 Cat marker 10 kb downstream P22 *att*B + TetA marker 29 kb upstream P22 *att*B | This work |
| JP21436 | JP18983 Cat marker 10 kb downstream P22 *att*B + TetA marker 32 kb upstream P22 *att*B | This work |
| JP21437 | JP18983 Cat marker 15.6 kb downstream P22 *att*B + TetA marker 29 kb upstream P22 *att*B | This work |
| JP21438 | JP18983 Cat marker 15.6 kb downstream P22 *att*B + TetA marker 32 kb upstream P22 *att*B | This work |
| JP21439 | JP18983 Cat marker 29 kb downstream P22 *att*B + TetA marker 32 kb upstream P22 *att*B | This work |
| JP20600 | JP20590 *prp*RΩTetA (47 kb downstream P22 *att*B, HF2) | This work |
| JP20601 | JP20591 *prp*RΩTetA (47 kb downstream P22 *att*B, HF2) | This work |
| JP20602 | JP20592 *prp*RΩTetA (47 kb downstream P22 *att*B, HF2) | This work |
| JP20605 | JP20595 *prp*RΩTetA (47 kb downstream P22 *att*B, HF2) | This work |
| JP20606 | JP20596 *prp*RΩTetA (47 kb downstream P22 *att*B, HF2) | This work |
| JP20607 | JP20597 *prp*RΩTetA (47 kb downstream P22 *att*B, HF2) | This work |
| JP20757 | JP20602 pJP2534 | This work |
| JP21704 | *E. coli* DC10B pJP2563 | This work |
| JP21747 | *E. coli* DC10B pJP2564 | This work |

| | | |
|---|---|---|
| JP21748 | *E. coli* DC10B pJP2565 | This work |
| JP21896 | *E. coli* DC10B pJP2566 | This work |
| JP21705 | *E. coli* DC10B pJP2567 | This work |
| JP21706 | *E. coli* DC10B pJP2568 | This work |
| JP21767 | VE18590 *celAΩtet*M (12.3 kb upstream of pp1 *att*B) | This work |
| JP21855 | VE18590 EF0370Ω*tetM* (12.3 kb downstream of pp1 *att*B, HF1) | This work |
| JP22112 | VE18590 EF0394Ω*tetM* (40.7 kb downstream of pp1 *att*B, HF2) | This work |
| JP22021 | VE18590 EF0480Ω*tetM* (121.1 kb downstream of pp1 *att*B, HF4) | This work |
| JP21771 | VE18590 EF2276Ω*tetM* (1.87 Mb downstream of pp1 *att*B) | This work |
| JP21768 | VE18562 *celAΩtetM* (12.3 kb upstream of pp1 *att*B) | This work |
| JP21856 | VE18562 EF0370Ω*tetM* (12.3 kb downstream of pp1 *att*B, HF1) | This work |
| JP21857 | VE18562 EF0394Ω*tetM* (40.7 kb downstream of pp1 *att*B, HF2) | This work |
| JP22110 | VE18562 EF0480Ω*tetM* (121.1 kb downstream of pp1 *att*B, HF4) | This work |
| JP21772 | VE18562 EF2276Ω*tetM* (1.87 Mb downstream of pp1 *att*B) | This work |

**Table S3. Oligonucleotides used in this study.**

| Mutagenesis | Primers | Sequence (5'-3') |
|---|---|---|
| **LT2 ∆Fels-1** | **LT2-AFels1-1m** | GGGGCACTCCTGGGGCAGTAGATGCCAGTTGTTGATTCAGTATATCTACTGTGTAGGCTGGAGCTGCTTCG |
| | **LT2-AFels1-2c** | GGCATCATACTGTACACTGTCATATGCCATATATTTAAACGCTAAAAGGGCATATGAATATCCTCCTTA |
| **LT2 ∆Gifsy-2** | **LT2-AGifsy2-13m** | CGCTGGAGTATACCTTGTTTAGCGATTTATTGAACCCCGATCACACCTGTGTAGGCTGGAGCTGCTTCG |
| | **LT2-AGifsy2-14c** | GCAAATCGCGCTACGCAGAATGTTCATCTTTTCAGGCACAAACGGCCGGTCCATATGAATATCCTCCTTAG |
| **LT2 ∆Gifsy-1** | **LT2-AGifsy1-1m** | CAAGTAACGAGGCGACATCAAACCTTGAGTTATTAAATAAATGGAGAATAGTGTGTAGGCTGGAGCTGCTTCG |
| | **LT2-AGifsy1-2c** | GTCCCTGATATGGGCGTGCAGGGTCGGTGAACTCCGTCAGGCTGAAGCATATGAATATCCTCCTTA |
| **LT2 ∆Fels-2** | **LT2-AFels2-1m** | CTTCAAGCTCCATGCTGGTCGTAAATCCATTATCCGGGCTGACAGAATGCTGTGTAGGCTGGAGCTGCTTCG |
| | **LT2-AFels2-2c** | TTCCTTGAGCATCTTCGGGCGGATCTCGGTAACGGGTGTGTTACCCAGAGCATATGAATATCCTCCTTA |
| **LT2 ∆*att*B P22** | **LT2-AattB-1m** | CCTTCTACCCCGTGATTCACCCGCGTGAACACACCCTTCTCATGTGTAGGCTGGAGCTGCTTCG |
| | **LT2-AattB-2c** | GGATACTGCTTATGTTTTGCTAGTTGTGTACCCGAGAGTGTACCCGGTCCATATGAATATCCTCCTTAG |
| **LT2 ∆*rfb*P-*rfb*B P22** | **LT2-ArfbP-1m** | TTTTACGCAGGCTAATTTATACAATTATTATTCAGTACTTCTCGGTAAGCTTGTGTAGGCTGGAGCTGCTTCG |
| | **LT2-ArfbB-2c** | TTTATTGGCAAATTAAATACCACATTAAATACGCCTTATGGAATAGAAAACATATGAATATCCTCCTTA |
| ***tet*A marker 6 kb upstream P22 *att*B** | **LT2-tetA-11m** | TCAGACACGATAAGTCTCCTTGGCGGTGGTCTGAAAAACGTTCTTACAGGCCACAGGGTTCGCTGTTAATCACTTTACTT |
| | **LT2-tetA-12c** | ACCCGCAGGCGGAGAAAGTGGTATTCTCAGTCGCATCTCGTAAGAGTTATCAATTCCGCTGGTTATCAAGAGGGTCATTA |
| ***tet*A marker 6 kb downstream P22 *att*B** | **LT2-tetA-3m** | GGTTGCGCATTATTCACGGCGGCCGTGGCTAAAACGCCCTGCATCCCCGCGCGGTTAAGGCGCTGTTAATCACTTTACTT |
| | **LT2-tetA-4c** | AATGCCGTAGCGTTTGCCACCGTTCACAAGATATTGAACCAGTTTCATTGCAGTCTTCTTGGTTATCAAGAGGGTCATTA |
| ***tet*A marker 47 kb downstream P22 *att*B** | **LT2-tetA-6m** | ATTGCATCATAAAAATACCCGCCTGGGTTTCCGGCGGGTATATTTATTCCTGGCAACCGGCGCTGTTAATCACTTTACTT |
| | **LT2-tetA-7c** | GCGTCCTGAATGCCTGATGGCGCTACGTTTACTGCAGGCCGTCATCCGGCAAAACGGATGGGTTATCAAGAGGGTCATTA |
| ***tet*A marker 88 kb downstream P22 *att*B** | **LT2-tetA-9m** | GCGAGGGAAGACTTAGTTTACCCGCGATTTCCGCGACGGTATGGATGAATCGTTGAGGCGCGCTGTTAATCACTTTACTT |
| | **LT2-tetA-10c** | CATTGATAAAAGGCTCGCGAATGCGAGCCTTTTTTATGCGTAAGGTGTTACGAACCACCTGGTTATCAAGAGGGTCATTA |
| ***tet*A marker 130 kb downstream P22 *att*B** | **LT2-tetA-13m** | CGTAAATATTGACTTGACATAGGCATCTACAGACCCGGCTTCTGCCGGGTCTTGTTGTTGCGCTGTTAATCACTTTACTT |
| | **LT2-tetA-14c** | GCTGATATGTCTCAGGACACCAGTATTCACGATTTTATCAAGCAAAACGCCCGTTAATAGGGTTATCAAGAGGGTCATTA |
| ***tet*A marker 170 kb downstream P22 *att*B** | **LT2-tetA-15m** | CTGCTGATTTTTATGTGGCGGATACCGGTGATTGATGAAAAGCAACAAAAGCGCTGAAGCCGCTGTTAATCACTTTACTT |
| | **LT2-tetA-16c** | TGTGCTCGAAAACGGTCGAATTTACTGGCTGTGAACGACAATTGCAACAGCGATTCGTGTGGTTATCAAGAGGGTCATTA |
| ***tet*A marker 213 kb downstream P22 *att*B** | **LT2-tetA-17m** | TCACTTTATATAAAAAGAAAGCAACGCAACGTATTGCTTCAGCTTAAAAATAATTATATCCGCTGTTAATCACTTTACTT |
| | **LT2-tetA-18c** | GATAACGTTGTTATGGGTTTAGCTATGAGGAACAAATAAATATAAAATAAAGAATTAGTAGGTTATCAAGAGGGTCATTA |
| ***tet*A marker 246 kb** | **LT2-tetA-19m** | TGTAGGAATCCCCGCCGCCCGTTACCCATTGGTGGCGGGGAACATTAATTATACATGAATCGCTGTTAATCACTTTACTT |

| | | |
|---|---|---|
| **downstream P22 *att*B** | **LT2-tetA-20c** | CATGGGACGTTAGAACAGGGGCGAGGACAAATGAGAATATTACGGAAATAATTAAATAACGGTTATCAAGAGGGTCATTA |
| ***tet*A marker 304 kb downstream P22 *att*B** | **LT2-tetA-25m** | GCCTTTGATTATCTGATTCGCACCAAAATCGCATAAAAGAAGTAAGCACACCTGCAAGGCGCTGTTAATCACTTTACTT |
| | **LT2-tetA-26c** | AAACACCGCAGGCCCGAATAGCTTACACTATCGGGCCATTTACGATGGCCAGTTAACTGGGGTTATCAAGAGGGTCATTA |
| ***tet*A marker 348 kb downstream P22 *att*B** | **LT2-tetA-27m** | GTGTGGCCACGATCAGTTCCAGCGCAGGCCGTTTGAGCCGTAAGGTTCATTATCGTGAGCCGCTGTTAATCACTTTACTT |
| | **LT2-tetA-28c** | GAAGAGATCCGCAAACTGGAAGCCCAACTGCACGCTTAACATGGCGGACGGCAACCGTCCGGTTATCAAGAGGGTCATTA |
| ***tet*A marker 392 kb downstream P22 *att*B** | **LT2-tetA-21m** | TGAGAAAGAAGCCGTTGAAATCGTTAGCGAAGTATTGAAAAACGCCTGATGGGCGATATGCGCTGTTAATCACTTTACTT |
| | **LT2-tetA-22c** | AGGCCTGATAAGCGCAGCGCCGTTAATACAAAAAAGGAGCCGTAAGGCTCCTTTTTCTTCGGTTATCAAGAGGGTCATTA |
| ***tet*A marker 425 kb downstream P22 *att*B** | **LT2-tetA-29m** | CGCATTGCCAGAAATAGCCGGAACCGACATCGGCGAGCGGCTATTGCCTGATGGCGCGACCGCTGTTAATCACTTTACTT |
| | **LT2-tetA-30c** | AGCAGTGGCCCGTTTTATATCAACCTATGCGCCCAAACAACCTTTGTAGGGCTGATAAGCGGTTATCAAGAGGGTCATTA |
| ***tet*A marker 474 kb downstream P22 *att*B** | **LT2-tetA-23m** | TATCGACCGTTATTATTATTCTTAATAAAAGGAGAGTGGTTCCAGAATGGCGCGCGGCGCCGCTGTTAATCACTTTACTT |
| | **LT2-tetA-24c** | AGGGCGAACGTTATAAGTACGTTTCCGGACGATGCAATAATTAAATGTATTATCAGAATGGGTTATCAAGAGGGTCATTA |
| ***cat* marker 6 kb downstream P22 *att*B** | **LT2-STM0331-cat-1m** | GGTTGCGCATTATTCACGGCGGCCGTGGCTAAAACGCCCTGCATCCCCGCGCGGTTAAGGGGCGCGCCTACCTGTGACGG |
| | **LT2-STM0331-cat-2c** | AATGCCGTAGCGTTTGCCACCGTTCACAAGATATTGAACCAGTTTCATTGCAGTCTTCTTGGAATAGGAACTTCATTTAAATGGC |
| ***tet*A marker 10 kb downstream P22 *att*B** | **LT2-stbE-tetA-2c** | CTCCTGAGTTTTGATGGGAAATATTCAGGGATACGATCGTATCCCTGAGGGGTTATCAAGAGGGTCATTA |
| | **LT2-stbE-tetA-1m** | ATCAGCTTAATGCTATTTTTGACAATGGTACATGCTGAGTCTGGCTTGCGCTGTTAATCACTTTACTT |
| ***tet*A marker 29 kb downstream P22 *att*B** | **LT2-STM0352-tetA-1m** | CGTTTCCCTCTTTACCGCAGCGTGTCGGCCATTCCGCAACGCTGCCGCAGCGCTGTTAATCACTTTACTT |
| | **LT2-STM0352-tetA-2c** | CCCGTCAGGAACGCTTGACCTTTCCTTCGTTGTAACGCCTAGCCTTTGCCGGTTATCAAGAGGGTCATTA |
| ***tet*A marker 32 kb downstream P22 *att*B** | **LT2-STM0355-tetA-1m** | CCGGTTTCCCGCCGCGCGAGAGGTAATCGTCAATGGCGATCGGTCTGGCGCGCTGTTAATCACTTTACTT |
| | **LT2-STM0355-tetA-2c** | CTCACGCAAAGAAAAGCGAAGCGTGGCTAGCGTATCGCGACCGGCCTGTGGTTATCAAGAGGGTCATTA |
| ***cat* marker 10 kb downstream P22 *att*B** | **LT2-stbE-cat-1m** | ATCAGCTTAATGCTATTTTTGACAATGGTACATGCTGAGTCTGGCTTGGGCGCGCCTACCTGTGACGG |
| | **LT2-stbE-cat-2c** | CTCCTGAGTTTTGATGGGAAATATTCAGGGATACGATCGTATCCCTGAGGGGAATAGGAACTTCATTTAAATGGC |
| ***cat* marker 15.6 kb downstream P22 *att*B** | **LT2-stbB-cat-1m** | CAACTCCCTAGCGATTGAAAATGGTCTGGCGTTTCAACGCCAGACCGTAGGGCGCGCCTACCTGTGACGG |
| | **LT2-stbB-cat-2c** | GGGACAGTCGCTGCCACCCTTCAATACGCCGTTTCGTATAAATAATTGTCGGAATAGGAACTTCATTTAAATGGC |
| ***cat* marker 29 kb downstream P22 *att*B** | **LT2-STM0352-cat-1m** | CGTTTCCCTCTTTACCGCAGCGTGTCGGCCATTCCGCAACGCTGCCGCAGGGCGCGCCTACCTGTGACGG |
| | **LT2-STM0352-cat-2c** | CCCGTCAGGAACGCTTGACCTTTCCTTCGTTGTAACGCCTAGCCTTTGCCGGAATAGGAACTTCATTTAAATGGC |
| **P22 Δ*int* (same primers used for ES18 Δ*int*)** | **P22-Aint-3m** | CCCGCTCGTTTTAATGCTGCCCTCCATGCAGTATTAGCGTCATAGTGTGTAGGCTGGAGCTGCTTCG |
| | **P22-Aint-4c** | GAGGAGAAGGCGCATAAGAAGTCGCTGGATGATGACAAGAGTCGGGGTCCATATGAATATCCTCCTTAG |

| | | |
|---|---|---|
| **P22 Δ*xis*** | **P22-Axis-1m** | ATGTCATCACCCGCGCTCACCTGGACAGTATGCAGCGGAG ATTGAAGTGCTGTGTAGGCTGGAGCTGCTTCG |
| | **P22-Axis-2c** | GAAACAGCGGAGTAAACATGGAATCACACAGCCTCACACTT GATGAGGCCGGTCCATATGAATATCCTCCTTAG |
| **P22 Δ*int-xis*** | **P22-Aint-3m** | CCCGCTCGTTTTAATGCTGCCCTCCATGCAGTATTAGCGTC ATAGTGTGTAGGCTGGAGCTGCTTCG |
| | **P22-Axis-2c** | GAAACAGCGGAGTAAACATGGAATCACACAGCCTCACACTT GATGAGGCCGGTCCATATGAATATCCTCCTTAG |
| **P22 Δ*orf12*** | **P22-Aorf12-1m** | GGTGGCTTGCTGATTGGCGGATTAACACCAACCGCCAGTG ACGTTCTGGCTGTGTAGGCTGGAGCTGCTTCG |
| | **P22-Aorf12-2c** | GCGGCTTCCTGATGGATGACAGGCGCTTTACAAGCTCGTC CATCGCTCTGGGTCCATATGAATATCCTCCTTAG |
| **ES18 Δ*xis*** | **ES18-Axis-1m** | TATGCCATCACCCGCGCTCACGGCGACAGTATGCATCGGA GACTGAAGAGTGTGTAGGCTGGAGCTGCTTCG |
| | **ES18-Axis-2c** | AGCAATAACAATCCTCGCACTCGCGGGGATTTCTTTTATCC GGAGTAACCGGTCCATATGAATATCCTCCTTAG |
| **ES18 Δ*int-xis*** | **P22-Aint-3m** | CCCGCTCGTTTTAATGCTGCCCTCCATGCAGTATTAGCGTC ATAGTGTGTAGGCTGGAGCTGCTTCG |
| | **ES18-Axis-2c** | AGCAATAACAATCCTCGCACTCGCGGGGATTTCTTTTATCC GGAGTAACCGGTCCATATGAATATCCTCCTTAG |

| **Plasmid** | **Primers** | **Sequence (5'-3')** | **Comment** |
|---|---|---|---|
| **pBAD18** | | | |
| **pJP2534** | **P22-int-5mS** | ACGCGTCGACCAAATACTTACGTATTATTCGTG CC | |
| | **P22-xis-4cE** | CCGGAATTCATTCTACGACATCGCTAACGC | |
| **pBT2bgal** | | | |
| **pJP2563** | **pp1-HF-1_FB** | AGTAGGATCCCAAAAGTAGGGGCCTTTTCTGC ACCTTTTACAGC | *celA* left flank |
| | **pp1-tetM_HF-1_R** | CTCAAATTGCGAGATTTGGGTTGCCTTTGTTTC TTGATCAAATTTGAACAAAGGACTGAAACGATA TGTCAATTTTGAAAAATG | |
| | **tetM-F** | GATCAAGAAACAAAGGCAACCCAAATCTCGCA ATTTGAG | *tetM* cassette |
| | **tetM-RS** | GTCAGTCGACGATCTTGTATCATTACTCCATGT ATCTATTGATG | |
| | **pp1-HF-1_FB** | AGTAGGATCCCAAAAGTAGGGGCCTTTTCTGC ACCTTTTACAGC | *celA* left flank-*tetM* fusion |
| | **tetM-RS** | GTCAGTCGACGATCTTGTATCATTACTCCATGT ATCTATTGATG | |
| | **pp1-HF-1_FS** | CATTTTTCAAAATTGACATATCGTTTCAGTCCTT TGTTCAAATTTGTCGACTTATACGTTTTGTGCTT CGACTTTTTCAAGGGTTTCTTC | *celA* right flank |
| | **pp1-HF-1_RH** | TGCAAAGCTTGTTTTCATTTTGTTTATGTCATCT GAAGTCAAAGATGCG | |
| **pJP2564** | **pp1-HF1_FB** | ATGAGGATCCGGCTACAACCAATTATTAGCTG ACGATTATGTG | EF0370 left flank |
| | **pp1-tetM_HF1-R** | CTCAAATTGCGAGATTTGGGTTGCCTTTGTTTC TTGATCTTAAATTGCTGTTAAATTAGGGTTATTG ATATTTTTAAAATAAAAATC | |
| | **tetM-F** | GATCAAGAAACAAAGGCAACCCAAATCTCGCA ATTTGAG | *tetM* cassette |
| | **tetM-RS** | GTCAGTCGACGATCTTGTATCATTACTCCATGT ATCTATTGATG | |
| | **pp1-HF1_FB** | ATGAGGATCCGGCTACAACCAATTATTAGCTG ACGATTATGTG | EF0370 left flank-*tetM* fusion |

| | | | |
|---|---|---|---|
| | **tetM-RS** | GTCAGTCGACGATCTTGTATCATTACTCCATGTATCTATTGATG | |
| | **pp1-HF1_FX** | AAATATCAATAACCCTAATTTAACAGCAATTTAATCTAGATTAAACAATCACTAAACAGAGGCTTGTCGACAG | EF0370 right flank |
| | **pp1-HF1_RNh** | GTCTGCTAGCTGGCGCAAAATGTTCATTATCTTGTGAAAAAAC | |
| **pJP2565** | **pp1-HF2_FB** | TGCAGGATCCACAACAATGCTAGATGCAGTTTTAGATGCGGACTC | EF0370 left flank |
| | **pp1-tetM_HF2-R** | CTCAAATTGCGAGATTTGGGTTGCCTTTGTTTCTTGATCTTAGGCTGAGTGTCCTACGATTGTGCTACCTGAGTAACC | |
| | **tetM-F** | GATCAAGAAACAAAGGCAACCCAAATCTCGCAATTTGAG | *tetM* cassette |
| | **tetM-RS** | GTCAGTCGACGATCTTGTATCATTACTCCATGTATCTATTGATG | |
| | **pp1-HF2_FB** | TGCAGGATCCACAACAATGCTAGATGCAGTTTTAGATGCGGACTC | EF0370 left flank-*tetM* fusion |
| | **tetM-RS** | GTCAGTCGACGATCTTGTATCATTACTCCATGTATCTATTGATG | |
| | **pp1-HF2_FS** | CTCAGGTAGCACAATCGTAGGACACTCAGCCTAAGTCGACTCAGCAATGATAAAAATAGACGTAAGCAGCTTC | EF0370 right flank |
| | **pp1-HF2_RNh** | ATGTGCTAGCATCATCAAATTGCAAATAACGGCAACCTCTCTC | |
| **pJP2566** | **pp1-HF4_FB** | ATGTGGATCCCATTTTAGTTAAATCAGTAATTTTCAATATCGAGGG | EF0480 left flank |
| | **pp1-HF4_RS** | GAAGCTATCTTAAAATTTAAAATTAACTAAGTCGACCTAAAAAAAGCCCCTTTCCACTTTTTCTG | |
| | **tetM-FS** | GTACGTCGACGATCAAGAAACAAAGGCAACCCAAATCTCGCAATTTGAG | *tetM* cassette |
| | **tetM-RS** | GTCAGTCGACGATCTTGTATCATTACTCCATGTATCTATTGATG | |
| | **pp1-HF4_FS** | CAGAAAAAGTGGAAAGGGGCTTTTTTTAGGTCGACTTAGTTAATTTTAAATTTTAAGATAGCTTC | EF0480 right flank |
| | **pp1-HF4_RH** | GTCTAAGCTTCTGATTAAAACTTTATTATGATGTCATAGTGAAG | |
| **pJP2568** | **2196839_FB** | ATGTGGATCCGAAATAAGCAATAATCATTGGTCAACGCCTCTC | EF2276 left flank |
| | **2196839_RS** | TAGATTATCGAATATTTTCGAAATTTAATTGAATGACAAGTCGACTCAAAGTACGAGGCGATACATTGCTGATAATTTTTTAGAGTATTG | |
| | **tetM-F** | GATCAAGAAACAAAGGCAACCCAAATCTCGCAATTTGAG | *tetM* cassette |
| | **tetM-RS** | GTCAGTCGACGATCTTGTATCATTACTCCATGTATCTATTGATG | |
| | **2196839_FB** | ATGTGGATCCGAAATAAGCAATAATCATTGGTCAACGCCTCTC | EF2276 left flank-*tetM* fusion |
| | **tetM-RS** | GTCAGTCGACGATCTTGTATCATTACTCCATGTATCTATTGATG | |
| | **2196839_FSa** | TGCAGTCGACTTGTCATTCAATTAAATTTCGAAAATATTCGATAATCTA | EF2276 right flank |
| | **2196839_RNh** | GTCAGCTAGCTCAGGTGTAGAAAATTTTACTGTAGATTTTGCC | |

**Table S4. Plasmids used in this study.**

| Plasmid | Description | Reference |
|---|---|---|
| **pKD46** | Amp$^{R}$. Thermosensitive plasmid with Red lambda system | (*31*) |
| **pCP20** | Amp$^{R}$. Thermosensitive plasmid with FLP recombinase | (*31*) |
| **pBAD18** | Amp$^{R}$. Expression vector | (*40*) |
| **pBT2βgal** | Amp$^{R}$, GN; Cm$^{R}$, GP. Vector for allelic replacement | (*41*) |
| **pRN6680** | pBluescriptΩ2.9-kb pMVN6 *Sma*I-*Hind*II [*tetA(M)*]; Amp$^{R}$, Tet$^{R}$ | (*42*) |
| **pJP2534** | pBAD18 *int-xis* P22 | This work |
| **pJP2563** | pBT2βgal with *tetM* from pRN6680 positioned between *E. faecalis* V583 flanking sequences at *celA*; Cm$^{R}$, Tet$^{R}$ | This work |
| **pJP2564** | pBT2βgal with *tetM* from pRN6680 positioned between *E. faecalis* V583 flanking sequences at *EF0370*; Cm$^{R}$, Tet$^{R}$ | This work |
| **pJP2565** | pBT2βgal with *tetM* from pRN6680 positioned between *E. faecalis* V583 flanking sequences at *EF0394*; Cm$^{R}$, Tet$^{R}$ | This work |
| **pJP2566** | pBT2βgal with *tetM* from pRN6680 positioned between *E. faecalis* V583 flanking sequences at *EF0480*; Cm$^{R}$, Tet$^{R}$ | This work |
| **pJP2568** | pBT2βgal with *tetM* from pRN6680 positioned between *E. faecalis* V583 flanking sequences at *EF2276*; Cm$^{R}$, Tet$^{R}$ | This work |
